# A curated human lactylome and protein language model framework enable accurate prediction and reveal local determinants of lysine lactylation

**DOI:** 10.64898/2026.07.28.741267

**Authors:** Zhengda Li, Youcheng Huang, Guangyao Shan, Di Zuo, Junhe Zhang, Yuqiang Du, Dejun Zeng, Xiliang Wang, Liang Chen, Hong Fan, Guangyu Yao

## Abstract

Lysine lactylation is a dynamic post-translational modification that can alter protein function and has been implicated in diverse physiological and pathological processes. Accurate identification of lactylation sites is therefore important for defining its regulatory landscape and for generating testable hypotheses about lactylation-associated mechanisms. Here, we introduce CLEAR-Lactyl and AttentionKla. CLEAR-Lactyl is a curated benchmark dataset of human lysine lactylation comprising 16,604 positive sites. AttentionKla is a deep learning framework trained on CLEAR-Lactyl that employs a pre-trained protein language model fine-tuned with LoRA; it significantly outperforms existing tools, and the factors contributing to its performance gain have been dissected through comprehensive ablation studies. Its utility in predicting novel lactylation sites and in sequence-directed modulation of lactylation levels has been experimentally validated in cellular assays. Together, CLEAR-Lactyl and AttentionKla provide a powerful platform for lysine lactylation research and offer an extensible framework for the precise modulation of other post-translational modifications.

## Introduction

Lysine lactylation (Kla) is a pan-species post-translational modification(PTM), where a lactyl group is conjugated to the ε-amino group of a lysine residue. This modification profoundly alters the physicochemical properties, structure, stability, and function of proteins, thereby playing key roles in diverse biological processes^1–3^. The lactylation process is governed by a dynamic equilibrium involving lactyltransferases, delactylases, lactyl-CoA synthetases, and multiple microenvironmental processes^4–6^. Reported lactyltransferases include p300, GCN5, and AARS1/2, canonical acetyltransferases and alanyl-tRNA synthetases that have also been reported to exhibit lactyltransferase activity^6,7^.

Conventional characterization of Kla sites relies on liquid chromatography-tandem mass spectrometry (LC-MS/MS), which identifies Kla sites with high throughput and accuracy by examining mass shifts of peptides. This technique remains the gold standard for discovering new sites, yet its sensitivity is limited, making it difficult to detect low-abundance sites or transient, dynamically modified sites, thereby constraining research on these modifications. A high-throughput, proteome-wide preliminary screening approach would substantially facilitate such studies. Deep learning models predict the probability of lactylation by learning sequence features around Kla sites, thus providing candidate screening for experiments. Existing lactylation prediction models include FSL-Kla, trained on human, mouse, and *Botryotinia fuckeliana* datasets based on few-shot learning plus ensemble deep learning; DeepKla, trained on a rice dataset using a CNN+BiGRU+Attention architecture; Auto-Kla, trained on a human gastric cancer cell line dataset using the AutoGluon framework; and PBertKla, trained on a human hepatocellular carcinoma tissue-derived dataset by fine-tuning ProteinBERT.^8–11However^, existing predictors were generally trained on relatively small and context-specific datasets, which may limit their robustness across broader proteomic settings and diverse species. Furthermore, these models were trained on small, species- or tissue-restricted datasets, limiting their applicability in large-scale proteome-wide analyses.More importantly, the ultimate value of precision predictive models lies not just in cataloging new sites, but in providing the foundational blueprint required for site-specific manipulation of Kla levels.

Accurate computational prediction is not simply a tool for discovery; it is a fundamental prerequisite for the rational, site-specific manipulation of Kla levels. As modulating Kla levels and controlling their functional outputs have become central focuses of lactylation research, current experimental strategies remain suboptimal. Approaches modelled after acetylation studies: mutating the positively charged lysine to an uncharged glutamine to mimic lactylation, or to arginine, which retains a positive charge but cannot be lactylated. These mutation-based methods can conveniently reveal certain mechanisms and roughly mimic the charge state of the lactylated/unlactylated residue; however, the spatial structure of the mutated residue differs from that of a genuine lactyllysine, making it impossible to determine whether the mechanistic alteration arises from the gain or loss of lactylation per se. Moreover, such mutations block other PTMs at the site, disrupting the dynamic and reversible PTM landscape at that position. A newly reported approach based on Genetic Code Expansion (GCE) constructs an orthogonal pyrrolysyl-tRNA synthetase (aaRS)/Klac-tRNA pair to directly and specifically incorporate lactyllysine (Klac) into a designated protein site. This strategy yields structurally authentic and site-specific lactylation, offers species universality, and holds potential for investigating dynamic Kla regulation.^12^ Nevertheless, this technique has a high technical barrier and a long application cycle. Currently, it can only achieve complete lactylation, masking other PTMs and dynamic PTM changes at that lysine residue.

Studies of phosphorylation and ubiquitination have shown that substitutions at residues flanking a modified site can alter local recognition motifs and thereby influence modification propensity^13–15^. Mutations within the substrate-recognition pockets of modifying enzymes can likewise alter substrate preference^16^. This suggests that the information governing PTM specificity is, to a significant extent, encoded in the local sequence context. Consistent with this, distinct amino acid preferences and motifs have been identified around Kla sites, including the strong enrichment of serine and proline at positions -1 and +3 around lactylation sites in human datasets, the less frequent enrichment of proline and glutamate at the +1 position, and the formation of specific motifs around lactylation sites in *Candida albicans*^12,17^. Therefore, definable recognition motifs also exist for lysine lactylation. However, PTM regulation is complex, involving multiple enzymes and microenvironmental factors, making isolated discoveries of modifying enzyme – substrate pairs insufficient. Through large-scale protein-sequence pre-training, protein language models such as ESM-2 learn contextual representations that capture evolutionary, structural, and functional constraints in protein sequences^18^. Inspecting the changes in model-estimated lactylation probability following flanking-residue substitutions may provide testable hypotheses regarding sequence determinants of lactylation. Hence, such substitutions could potentially alter the lactylation propensity of the central lysine while preserving the lysine residue itself, which minimizes the disruption to the central lysine’s PTM landscape.

Here, we present CLEAR-Lactyl, a comprehensive and meticulously curated benchmark dataset of human lysine lactylation. We leverage this dataset to fine-tune the large protein language model ESM-2, yielding AttentionKla, an enzyme-agnostic predictor that captures high-dimensional sequence features surrounding Kla sites. Through in silico saturation mutagenesis, AttentionKla reveals a previously underappreciated contribution of local sequence context to Kla regulation. Guided by these predictions, we designed and experimentally validated single-point mutations in flanking residues that lead to rational, site-specific modulation of Kla levels, independent of any modifying enzyme’s known substrate motif. This work establishes a novel and broadly applicable framework for predicting and precisely manipulating lysine lactylation, opening new avenues for functional interrogation of this critical PTM.

CLEAR-Lactyl, the independent test sets, AttentionKla source code and model weights, and proteome-wide predictions for human lysine sites are all publicly available. This also provides a website for online prediction using the AttentionKla model.

## Results

### An overview of the study

In this study, we present a deep-learning framework for the accurate prediction and mechanistic exploration of protein lactylation (Figure 1). To overcome the limitations of existing small-scale and single-source datasets, we constructed CLEAR-Lactyl, a large-scale, high-quality, homology-reduced dataset for human lactylation. Notably, our data processing pipeline inherently accommodates truncated sequences using padding strategies, preserving critical terminal signals.

**Figure 1.**
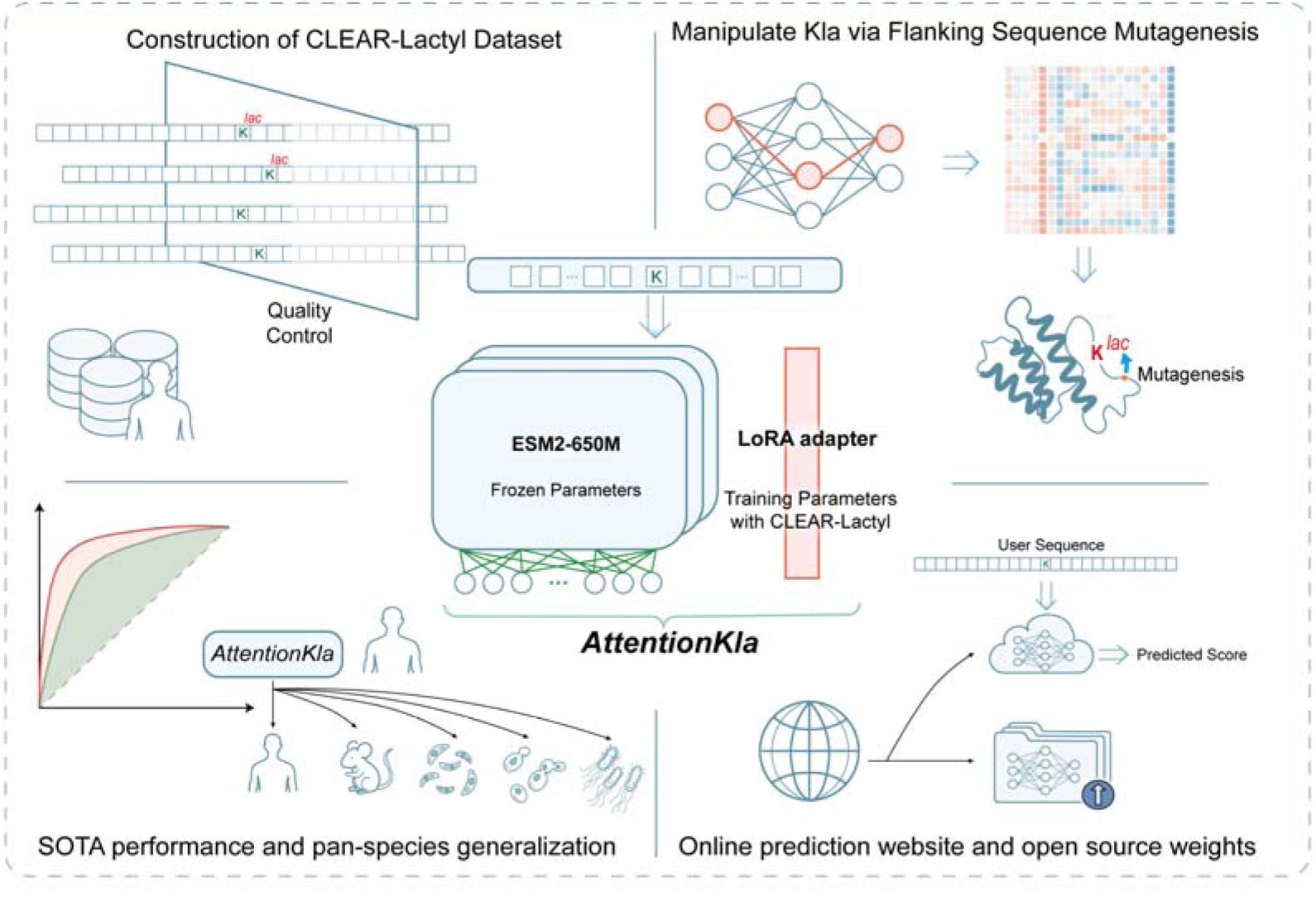
Overview of the AttentionKla framework and study design. The schematic illustrates the systematic workflow encompassing data construction, model development, performance evaluation, mechanistic application, and open-source deployment. **Construction of CLEAR-Lactyl Dataset**: A large-scale, high-quality dataset was curated through stringent quality control, homology reduction, and appropriate handling of truncated sequences using <PAD> tokens. **Model Architecture**: AttentionKla is built upon the ESM2-650M pre-trained foundation model. The core parameters are frozen, and lactylation-specific training is achieved efficiently via a Low-Rank Adaptation (LoRA) adapter using the CLEAR-Lactyl dataset. **SOTA Performance and Pan-species Generalization**: AttentionKla achieves superior predictive performance (ROC curves) and demonstrates robust generalization across multiple species, including human, mouse, and various microorganisms. **Manipulate Kla via Flanking Sequence Mutagenesis**: By extracting features from the neural network and analyzing the attention heatmap, critical flanking residues are identified. In silico predictions guide targeted mutagenesis on the protein structure to artificially manipulate specific lactylation (Kla) levels. **Online Prediction and Open Source**: The trained model is deployed as a user-friendly online web server, and the model weights are open-sourced to support customizable batch predictions for customer sequences.

At the core of our framework is AttentionKla, a lightweight yet highly robust predictor built upon the pre-trained protein language foundation model, ESM2-650M. By freezing the foundation model parameters and employing a Low-Rank Adaptation (LoRA) adapter, AttentionKla effectively learns lactylation-specific features with optimal computational efficiency. Comprehensive benchmarking demonstrated that AttentionKla achieves state-of-the-art (SOTA) performance in predicting human lactylation sites and exhibits remarkable pan-species generalization capabilities across evolutionary diverse organisms.

Beyond sequence-level prediction, we integrated attention-based interpretation with in silico mutagenesis to investigate the sequence determinants of protein lactylation. Position-specific attention profiles were first used to identify flanking residues prioritized by the model, and selected residue substitutions were subsequently evaluated were subsequently evaluated to predict their potential effects on lactylation propensity. These analyses guided the selection of candidate residues for experimental validation, through which targeted mutations were shown to modulate the lactylation levels of specific sites. This attention-guided computational and experimental framework connects model interpretability with the identification and manipulation of biologically relevant sequence determinants. To broaden the accessibility and utility of AttentionKla, we further developed a publicly available web server and released the trained model weights, enabling users to perform customized predictions on user-provided protein sequences.

### Construction of the CLEAR-Lactyl dataset and characterization of lactylation sequence features

Although a growing body of work has investigated the functional consequences of lactylation, the public availability of raw MS data from lactylation-antibody enrichment experiments remains limited. To circumvent this limitation, we compiled MS-derived human lactylation-site data from published studies^19–31,7,12,32,3^. (Figure 2A) These datasets were derived from both tumor and non-tumor tissues/cells(detailed information is provided in the *Source Data*). Analysis of the 5 amino acids flanking the lactylation sites across various datasets revealed shared surrounding sequence motifs, specifically characterized by lysine being the most frequently occurring amino acid in these flanking regions (Figure 2B).To mitigate potential data leakage caused by sequence homology, the positive samples were subjected to de-duplication and homology reduction using CD-HIT^33^, yielding 16,604 unique lactylation sites across 4,934 proteins. Overall, an elevated proportion of lysine residues was also observed at positions -5 to -20 and +5 to +20 in the flanking regions of lactylation sites (Figure 2C). At the protein level, the proportion of lactylated lysines varied across different proteins, with 47.6% of the proteins exhibiting lactylation on less than 10% of their lysine residues (Figure 2D). Spatially, these lactylation sites tended to be localized within the central regions of the protein sequences. However, 5% of the sites were distributed within the first 6.4% or the last 6.7% of the sequences. This distribution may partly reflect the limited detectability of N- and C-terminal peptides by MS-based workflows. Alternatively, lactylation may be intrinsically less frequent near protein termini(Figure 2E).

**Figure 2.**
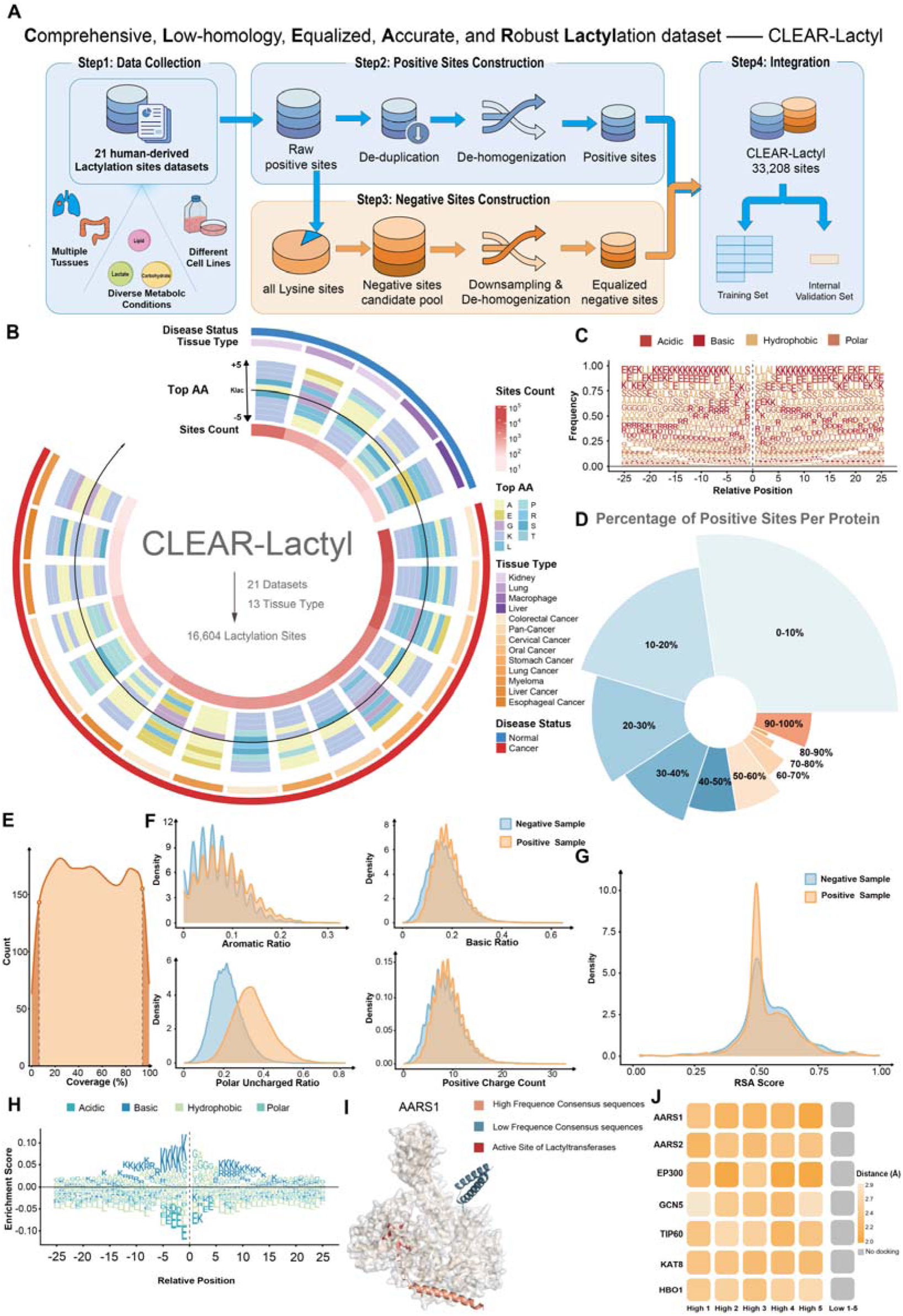
Construction of the CLEAR-Lactyl dataset and characterization of lactylation site features. (A) Schematic overview of the CLEAR-Lactyl dataset construction pipeline. **(B)** Sequence motif analysis of the 5 amino acids flanking the lactylation sites across different datasets, highlighting the enrichment of lysine residues. **(C)** The proportion of lysine residues at remote flanking positions (-5 to -20 and +5 to +20) relative to the lactylation sites. **(D)** Distribution of the percentage of lactylated lysines within individual proteins. **(E)** Spatial distribution of lactylation modification sites across the full length of protein sequences, showing the positional variation at the N- and C-termini. **(F-G)** Comparison of physical and chemical sequence features, including aromatic ratio, basic ratio, polar uncharged ratio, and positive charge count, between positive (lactylated) and negative (unlactylated) sequences. **(H)** Distribution plot illustrating the differences in amino acid frequencies (positive minus negative sequences), demonstrating the characteristic increase in lysine frequency. **(I-J)** AlphaFold3-predicted interaction models comparing the binding capacities of the highest- and lowest-frequency consensus sequences to the active pockets of lactylation enzymes.

Furthermore, lysine residues without detected lactylation within the same proteins containing positive sites were utilized to construct a negative data candidate pool. These “hard samples” inherently reflect the sequence characteristics of unlactylated sites. Following homology reduction, to maintain class balance, an equal number of negative samples was randomly selected from the candidate pool, resulting in a final negative dataset comprising 16,604 sites across 3,944 proteins. Collectively, these data construct CLEAR-Lactyl, encompassing a total of 33,208 site entries. **CLEAR-Lactyl** stands for a **C**omprehensive, **L**ow-homology, **E**qualized, and **A**ccurate dataset for **R**obust **Lactyl**ation prediction. Given that CLEAR-Lactyl currently represents the largest positive site dataset available, true positive sites could be more rigorously excluded during the generation of the negative candidate pool. This endows the randomly selected negative samples with a lower false-negative rate, a reduction that is anticipated to contribute significantly to the improvement of overall model performance.

Compared to negative sequences, the sequences containing lactylation sites exhibited an increased distribution in the aromatic ratio, basic ratio, proportion of polar uncharged amino acids, and positive charge count, whereas the Relative Solvent Accessibility (RSA) scores showed no significant changes (Figures 2F-G, Supplementary Figure 1A). The distribution plot of amino acid frequencies in positive sequences minus those in negative sequences further highlighted a characteristic enrichment of lysine residues (Figure 2H). This discrepancy may be associated with the substrate recognition mechanisms and charge neutralization features inherent to lactylation modifications.

Furthermore, motif discovery performed on the sequences encompassing flanking amino acids of the positive sites predominantly identified lysine-formed motifs (Supplementary Figures 1B-C). These results suggest that the lysine-enriched motifs flanking lactylation sites play a crucial role in the lactylation process. The underlying mechanism may involve a higher binding affinity between the positively charged lysines and the acidic pockets of lactylation enzymes (writers).

To verify whether the amino acids flanking the target lysine influence its binding capacity to lactylation enzymes, we utilized AlphaFold3 to predict the complex structures of these enzymes paired with consensus peptides constructed from the highest and lowest frequency amino acids found in the positive sequences. Strikingly, we observed that the top 5 highest-frequency sequences successfully interacted with the active pockets of 7 different lactylation enzymes. In contrast, the 5 lowest-frequency sequences failed to bind to these active pockets. (Figures 2I-J, Supplementary Figure 1D) Notably, this binding pattern was consistent regardless of whether the lactylation enzyme was an acetyltransferase or an alanyl-tRNA synthetase. Structural modeling is consistent with the possibility that flanking sequence context contributes to lactylation enzymes recognition.Taken together, these findings indicate that the enrichment of lysine residues near the modification sites may enhance the substrate recognition capacity of lactylation enzymes through mechanisms such as electrostatic interactions, thereby leading to elevated levels of lactylation.

### Functional landscape of the human lactylome reveals extensive involvement in metabolism and post-transcriptional regulation

To comprehensively characterize the biological functions of human protein lactylation, we performed systematic enrichment analyses on the 4,934 lactylated proteins from the CLEAR-Lactyl dataset. KEGG pathway enrichment revealed that lactylation modifications are not only highly concentrated in classical carbon and energy metabolism pathways (including glycolysis/ gluconeogenesis, the citrate cycle, and pyruvate metabolism), but also significantly enriched in core complexes essential for maintaining cellular homeostasis, such as the spliceosome, ribosome, and proteasome. (Supplementary Figure 2A) Interestingly, pathways associated with various neurodegenerative diseases (e.g., Parkinson’s disease and Huntington’s disease) also displayed significant enrichment, indicating a potential role for lactylation in neuropathological processes.

Further Gene Ontology (GO) analysis corroborated and expanded upon these findings (Supplementary Figure 2B). In terms of Molecular Function (MF), a substantial number of lactylated proteins exhibited RNA binding and oxidoreductase activities. Cellular Component (CC) enrichment analysis highlighted annotations related to the nuclear lumen, mitochondrial matrix, and focal adhesions. The significantly enriched terms for Biological Processes (BP) were overwhelmingly dominated by RNA metabolism, including RNA splicing, mRNA processing, and ribosome biogenesis.

By constructing a functional enrichment network (Supplementary Figure 2C), we identified several highly interactive modules regulated by lactylation: a post-transcriptional regulation hub centered on “RNA splicing and processing,” a protein synthesis hub centered on “ribosome assembly and translation,” as well as functional clusters associated with “DNA replication and repair” and “fatty acid oxidation.” Collectively, these results reveal that lactylation modifications are extensively and deeply integrated into the regulatory networks of cellular metabolism and post-transcriptional gene control.

### Development of AttentionKla for accurate, whole-proteome, and cross-species lactylation prediction based on protein language models

AttentionKla was trained using CLEAR-Lactyl, with 10% of the dataset reserved as an internal test set and the remaining 90% used for training and validation. AttentionKla is a model fine-tuned from the ESM2-650M pre-trained protein language model using Low-Rank Adaptation (LoRA). Input sequences consisted of 51 amino acids (aa) centered around the target lysine (25 aa upstream and downstream). Approximately **12.8%** of the lactylated lysine sites in CLEAR-Lactyl are located near the N- or C-terminus of proteins, resulting in truncated 51-aa windows with missing terminal residues. Although these sites account for a minor proportion, the terminal regions of proteins are rich in functional signals; thus, incorporating them into the training process is crucial for the model’s practical utility. To address this, we utilized the <PAD> token of the foundational ESM2 model to fill in the missing terminal positions, ensuring the model could process the full-length sequence windows. We allocated 1/10 of the total data as an internal testing set and used the remaining data for 9-fold cross-validation training, strictly ensuring that no identical site sequence appeared across different subsets to prevent data leakage. Hyperparameters were optimized using Optuna^34^ (Supplementary Figure 3A-B). Using the optimal hyperparameter combination, the ESM2 model was fine-tuned for 13 epochs until convergence, culminating in the AttentionKla model. To understand the model’s representation capabilities, we extracted sequence features from different layers of AttentionKla and reduced their dimensionality into a UMAP space (Supplementary Figure 3C). The results demonstrated that as the site data progressed through the model’s layers, AttentionKla effectively extracted lactylation-related features and successfully separated positive from negative sites.

We selected DeepKla, AutoKla, and PBertKla as existing published predictors for performance comparison. Given that DeepKla and PBertKla cannot predict terminally truncated sequences, this round of benchmarking only included AutoKla. On the internal testing set, AttentionKla achieved an AUC of 0.86(Figure 3A)(Other performance indicators are reported in Source Data). Benefiting from the expanded training set and the fine-tuning of the pre-trained protein model, its performance was significantly improved compared to existing methods. Moreover, the strict leakage-free data splitting and the large-scale testing set make the results highly representative of real-world application scenarios.Furthermore, we constructed an independent external testing set originating from disease profiles and tissues distinct from CLEAR-Lactyl. The positive sites were derived from human placental choriocarcinoma cell line data and were de-duplicated against the CLEAR-Lactyl dataset^35^. Since the external dataset provided only positive sites, we sourced the negative sites from the unused candidate pool of CLEAR-Lactyl. This specific selection strategy was adopted because the CLEAR-Lactyl candidate pool inherently contains fewer false negatives, allowing for a more reasonable and accurate evaluation of model performance without interference from false-negative noise. This process yielded a final external testing set comprising 4,426 site entries. On this external set, AttentionKla achieved an AUC of 0.89 (Figures 3B-C), demonstrating robust generalization capabilities across data from different human diseases and tissue sources. Notably, the external testing AUC was slightly higher than the internal testing AUC; we speculate this is because homology reduction was not applied to the external set, leading to a more lenient evaluation. To ensure a fair benchmark comparison with DeepKla and PBertKla, which cannot process truncated sequences, we removed all site sequences requiring <PAD> filling from the CLEAR-Lactyl dataset and re-trained a version without padding, named AttentionKla-nopad. On the corresponding padding-free internal and external testing sets, AttentionKla-nopad achieved AUCs of 0.88 and 0.86, respectively (Figures 3D-F), still demonstrating substantial performance improvements over DeepKla and PBertKla.

**Figure 3.**
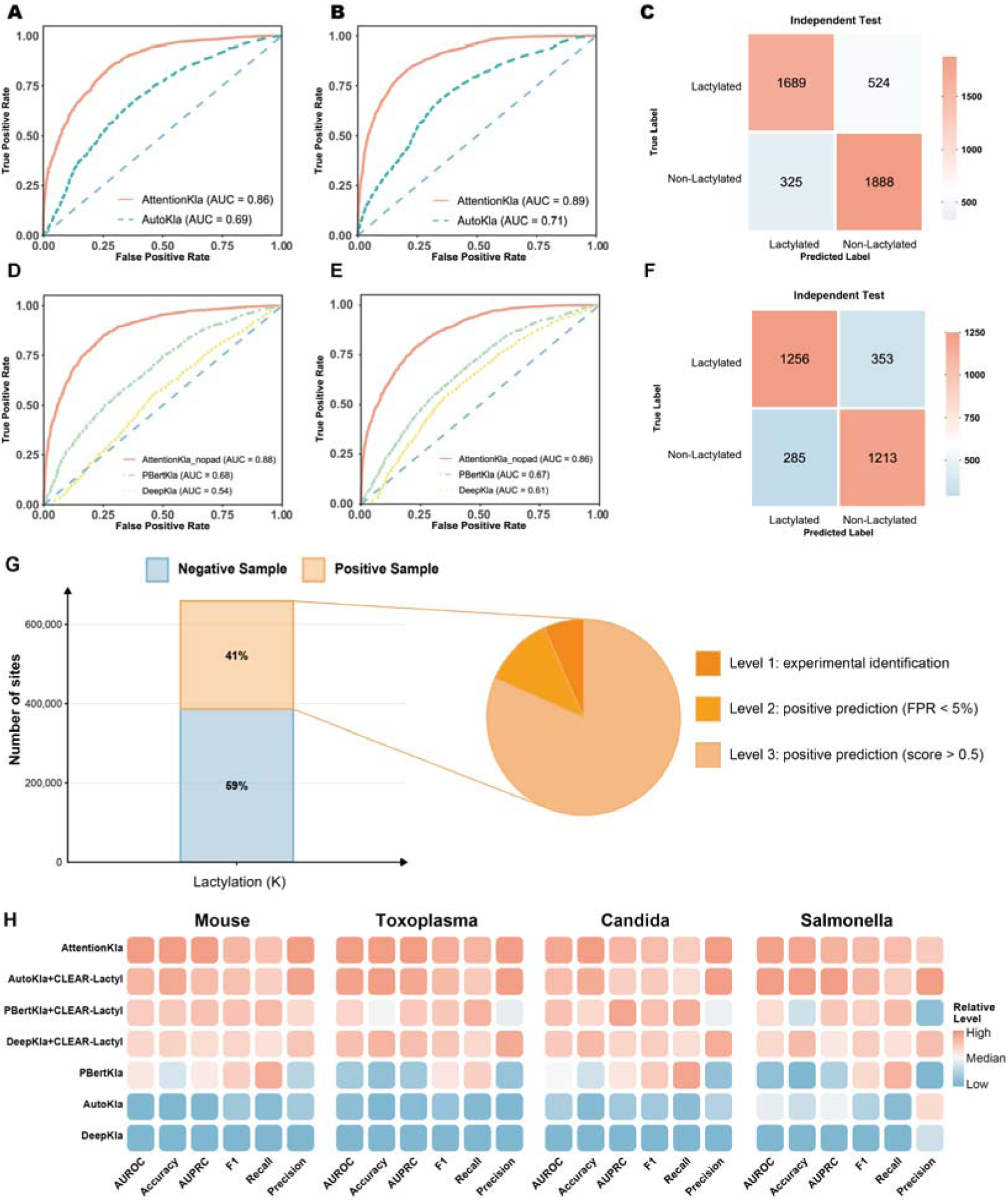
Performance evaluation and generalization capability of AttentionKla. **(A)** ROC curves and accuracy (ACC) of AttentionKla compared to AutoKla on the internal testing set. **(B-C)** Performance evaluation on the external independent testing set (derived from human placental choriocarcinoma cells), demonstrating model generalization across different tissue and disease contexts. **(D-F)** Benchmark comparisons of the AttentionKla-nopad version against DeepKla and PBertKla on both internal and external datasets, excluding padding-required truncated sequences. **(G)** Whole-proteome scan results predicting lactylation potential across all human lysines in the UniProt database, highlighting the proportion of high-confidence sites (FPR < 5%). *(Note: H and I are currently missing in the text)***(H)** Cross-species generalization performance of AttentionKla compared with baseline models across four distinct species (*Mouse, Toxoplasma, Candida, Salmonella*), including evaluations with and without baseline re-training on the CLEAR-Lactyl dataset.

Given AttentionKla’s robust predictive power and its capability to evaluate terminally truncated sequences, we deployed the model to scan and predict lactylation potentials across all human protein lysine sites. The results indicated that approximately 41% of lysine sites possess lactylation potential. Among these predicted positive sites, high-confidence sites with a false positive rate (FPR) of less than 5% accounted for 11.74% (Figure 3G). Although the experimentally observed abundance of lactylation modifications is currently relatively low, our estimate is consistent with the predictions of published pan-PTM models for other types of lysine PTMs, in which the estimated proportions of potential lysine sites varied across modification types, including approximately 40% for acetylation, 56% for methylation, 45% for SUMOylation, and 45% for ubiquitination. These estimates place the predicted lactylation proportion within the range reported by analogous sequence-based pan-PTM models, but do not constitute experimental validation^36^.

AttentionKla was fine-tuned on the pan-species pre-trained protein model ESM2-650M, suggesting it may possess cross-species lactylation site recognition capabilities. To verify this, we collected lactylation modification site data from four species with varying evolutionary distances from humans—*Mouse*, *Toxoplasma*, *Candida*, and *Salmonella*^17,37–39^—to serve as the cross-species positive set. We constructed a cross-species generalization testing set by pairing these with an equal number of unreported lysine sites from the same species as negative controls. Although the limited number of positive sites inevitably meant the negative set might contain potential false negatives, AttentionKla outperformed existing tools across all four species (Figure 3H). Even when comparator models such as DeepKla were re-trained using the CLEAR-Lactyl dataset to enhance their performance, AttentionKla maintained its leading position, reflecting the advanced nature of its architecture. Notably, PBertKla, which was also fine-tuned from a pan-species pre-trained model (ProteinBert), exhibited higher cross-species performance than the non-pan-species pre-trained AutoKla and DeepKla. This further suggests that fine-tuning strategies based on pan-species pre-trained models can enhance a model’s generalized prediction capability for pan-species lactylation sites.

### Ablation analyses define the contributions of model design, training-data scale, and terminal-sequence padding to AttentionKla performance

To determine the specific contributions of different modules to the performance improvement of AttentionKla, we performed comprehensive ablation experiments on both the model architecture and the CLEAR-Lactyl dataset. The AttentionKla model (ESM2-650M fine-tuned with LoRA) exhibited a significant performance improvement compared to fine-tuning only the last layer of ESM2-650M. Meanwhile, LoRA fine-tuning achieved performance comparable to full-parameter fine-tuning on the internal testing set, while additionally gaining performance improvements on the external testing set (Figure 4A, Supplementary Figure 4A). LoRA fine-tuning not only mitigates potential overfitting risks, but from a deployment perspective, its weights can be linearly merged into the original foundation model weights without introducing any inference latency. It requires storing only a minimal task-specific adapter file, facilitating lightweight switching among multiple tasks on the same foundational model. Consequently, LoRA represents the optimal strategy in this study for balancing performance and computational efficiency.

**Figure 4.**
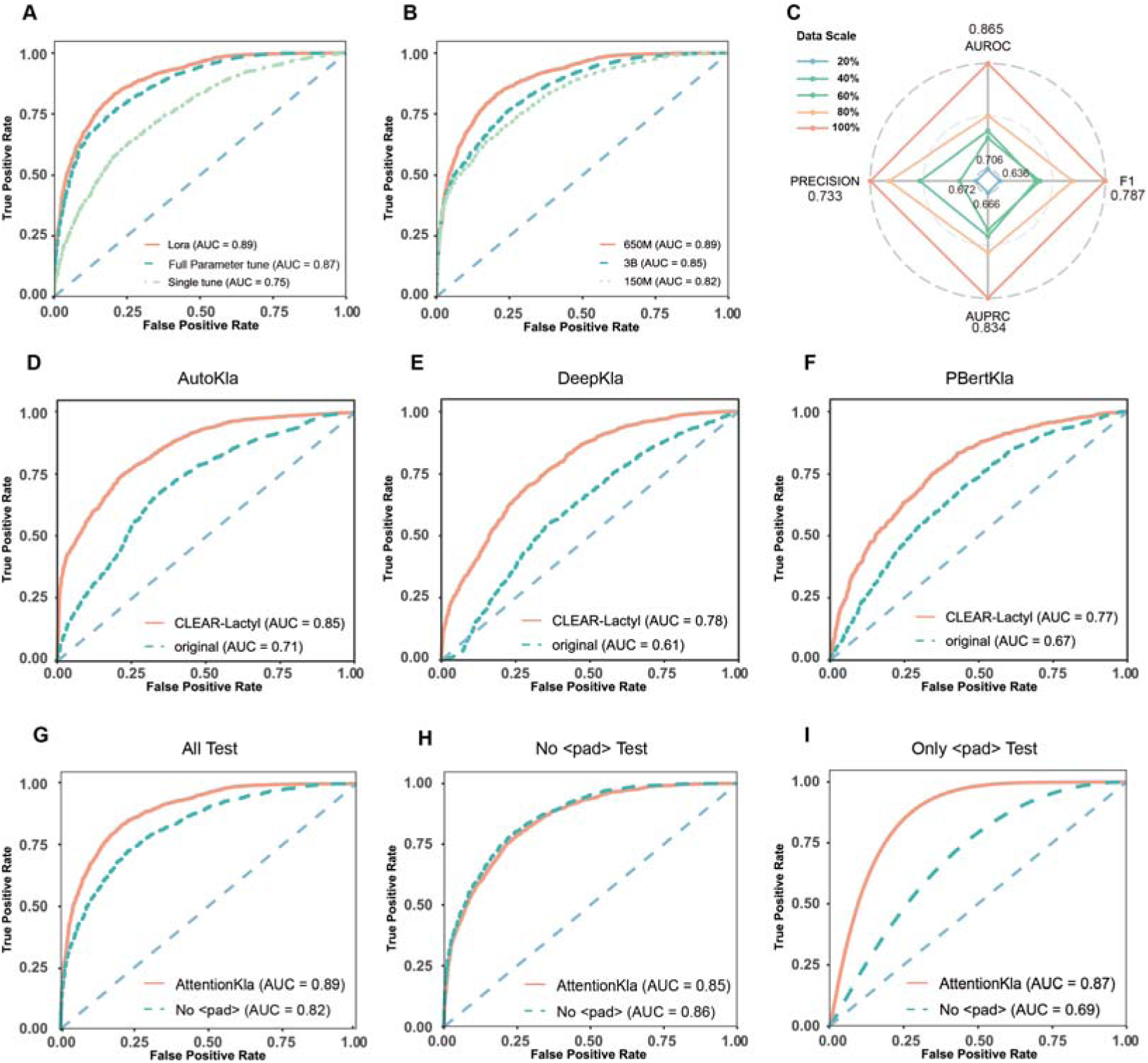
Ablation studies and performance optimization of AttentionKla on the external set. **(A)** Performance comparison of different fine-tuning strategies (LoRA, full-parameter, and last-layer fine-tuning) evaluated on the external set. **(B)** Impact of foundation model parameter scales (ESM2-650M vs. 3B) on the performance of AttentionKla on the external set. **(C)** Dataset scaling law analysis showing model performance variation when trained on randomly subsampled subsets of CLEAR-Lactyl, evaluated on the external set. **(D-F)** Performance improvements (AUC increments) of baseline models (AutoKla, DeepKla, and PBertKla) evaluated on the external set after being re-trained with the CLEAR-Lactyl dataset. **(G-I)** Performance comparisons between AttentionKla and AttentionKla-nopad on various external sets: **(G)** mixed sets containing both <PAD> and no <PAD> sequences, **(H)** strictly only-no <PAD> sequences, and **(I)** strictly only-<PAD> sequences.

To investigate the impact of foundation model parameter scales, we compared the performance of ESM2 models with different parameter sizes fine-tuned via LoRA. We determined that the performance of AttentionKla-650M was comparable to that of AttentionKla-3B, successfully balancing predictive performance with deployment feasibility (Figure 4B, Supplementary Figure 4B).

Because Kla is unlikely to occur at a 1:1 ratio of modified to unmodified lysine sites in vivo, the use of a balanced training set could raise concern about false positives when the model is applied to more imbalanced settings. To test whether AttentionKla learned sequence determinants beyond the training-set class prior, we constructed a strictly non-overlapping external imbalanced test set containing 1,391 positive sites and three negative sites per positive site (1:3). A matched balanced subset was generated by retaining all positive sites and randomly sampling one third of the negatives. AttentionKla achieved an AUROC of 0.88 in both settings, and the MCC changed minimally after increasing the number of negatives (0.6256 vs. 0.6521; Supplementary Figure S4C), indicating robust discrimination under this prevalence shift.

Given that the number of sites in CLEAR-Lactyl is substantially larger than the training sets of existing models, we investigating whether a scaling law exists regarding dataset size. By randomly subsampling the existing training set to create datasets of varying scales, we observed a significant decline in model performance as the data size decreased (Figure 4C). This suggests that even larger datasets hold the potential to further enhance the prediction of lactylation modifications. Furthermore, when baseline models (AutoKla, DeepKla, and PBertKla) were re-trained using CLEAR-Lactyl (with some utilizing the no <PAD> version), their AUCs increased by 0.14, 0.22, and 0.09 in external dataset, respectively (Figures 4D-F, Supplementary Figures 4C-E). Despite this data-driven enhancement, the performance of these baseline models still fell short of AttentionKla, underscoring the architectural superiority of the ESM2+LoRA fine-tuning approach.

To evaluate the impact of including terminally truncated sequence data filled with <PAD> tokens, we compared the performance of the AttentionKla and AttentionKla-nopad models across different testing sets. On testing sets containing both <PAD> and no <PAD> sequences, AttentionKla exhibited a significant performance improvement over AttentionKla-nopad, indicating that incorporating <PAD> data substantially elevates the overall capacity for lactylation prediction (Figure 4G, Supplementary Figure 4F). Specifically, when evaluating strictly on the no <PAD> testing sets, AttentionKla did not show a noticeable performance drop compared to AttentionKla-nopad. This indicates that training with <PAD>-inclusive data does not compromise AttentionKla’s performance on no-<PAD> sequences (Figure 4H, Supplementary Figure 4G).

Unsurprisingly, on the “only-<PAD>” testing set, AttentionKla vastly outperformed AttentionKla-nopad, providing a powerful predictive capability for truncated sequences (Figure 4I, Supplementary Figure 4H). These results indicate that including padded terminal windows during training is important for recovering lactylation sites near protein termini that would otherwise be excluded from fixed-window predictors. Notably, within the “only-<PAD>” testing set we observed that although <PAD> tokens occupy positions that would otherwise contain encoded amino-acid residues, they provide explicit terminal-boundary information. Consequently, the reduced residue content within the window did not substantially compromise prediction of terminally truncated sequences.. This may be attributed to the fact that <PAD> tokens can effectively provide implicit positional context—such as proximity to the protein termini—analogous to the functional information conveyed by encoded amino acids.

### AttentionKla guides the discovery of novel lactylation sites and precise modulation of lactylation levels via attention mechanisms

Although the performance of AttentionKla provides robust predictive support for wet-lab experiments, it is crucial to demonstrate how to integrate this model into existing experimental workflows to yield novel biological insights. Taking p53 as an example, previous studies have reported that lactylation at K120 and K139 affects its DNA-binding capacity^12^. Meanwhile, post-translational modifications (PTMs) in other domains of p53 are associated with tetramerization and nuclear localization^40^. Therefore, identifying novel lactylation sites may provide new insights into the functional regulation of p53. We performed inference on all lysine sites of p53 using AttentionKla and selected four “predicted positive” sites (K292, K321, K370, and K373) to construct arginine (R) mutants. Subsequent p53-immunoprecipitation (IP) assays confirmed that the K370R mutation resulted in a decreased lactylation level similar to that of the known K120R mutation.The reduced lactylation signal observed for K370R was consistent with a contribution of K370 to p53 lactylation and warrants site-specific validation (Figure 5A). Following transfection of HA-P53 into HEK293T cells, lactylated P53 was enriched using a pan-lactyllysine antibody, and targeted LC-MS/MS analysis identified lactylation at K370(Supplyment Data). Together with the reduced pan-Kla signal caused by the K370R mutation, these data support K370 as a lactylated residue of P53.This highlights the capability of AttentionKla in assisting the discovery of novel lactylation sites.

**Figure 5.**
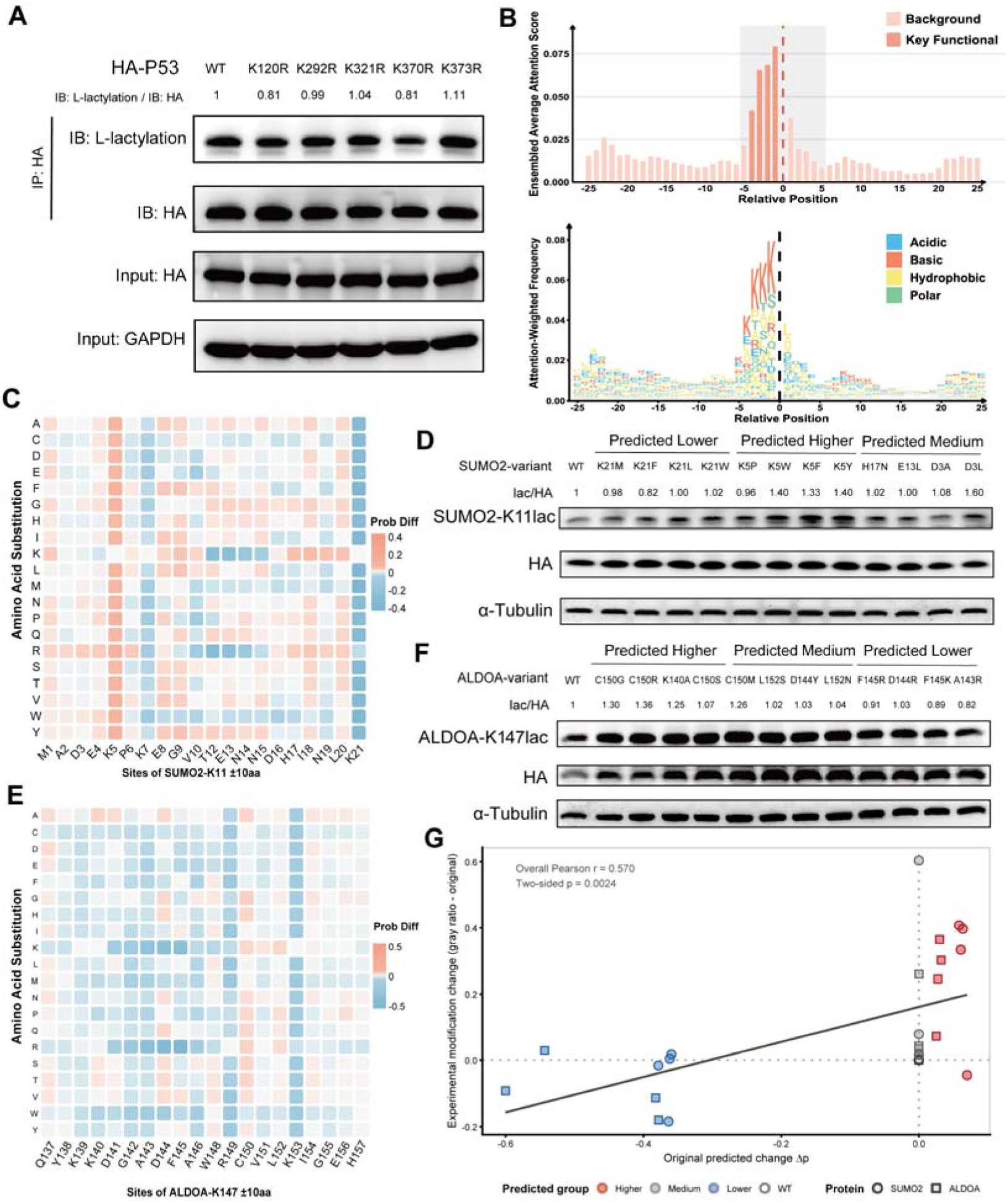
Experimental validation of AttentionKla for novel-site discovery and prediction-guided modulation of lysine lactylation. **(A)** Immunoprecipitation (IP) and immunoblot analyses identify p53-K370 as a candidate lactylation site. The K370R mutation decreased p53 lactylation, consistent with the K120R positive-control mutation; additional predicted candidates (K292R, K321R, and K373R) are shown. Lactylation signals were normalized to HA-tagged p53. **(B)** Schematic of the attention-extraction strategy. Integrated mean attention scores of the [CLS] token across sequence positions were used to identify residues that contribute to lactylation-site prediction. **(C,E)** In silico mutagenesis maps for flanking residues surrounding SUMO2-K11la (C) and ALDOA-K147la (E). Each cell represents the predicted change in lactylation probability (Δp) after substitution of the indicated amino acid at the corresponding flanking position. **(D,F)** Experimental validation of prediction-guided mutations at SUMO2-K11la (D) and ALDOA-K147la (F) using site-specific lactylation antibodies. For each site, variants were selected from mutations predicted to decrease, increase, or minimally affect lactylation, and the corresponding lactylation/HA signals are shown above the blots. **(G)** Association between model-predicted Δp and experimentally measured changes in site-specific lactylation across all variants. Point color denotes the predicted group (Higher, Medium, Lower, or WT), and point shape denotes the protein (SUMO2 or ALDOA). Model predictions were positively correlated with experimental changes for SUMO2-K11la (Pearson r = 0.531, P = 0.0617, n = 13) and ALDOA-K147la (Pearson r = 0.646, P = 0.0172, n = 13). The pooled analysis was also significant (Pearson r = 0.570, P = 0.0024, n = 26).

Across 33 TCGA cancer types, 84,943 of 728,821 missense SNVs (11.65%) were predicted to alter lysine lactylation. In absolute numbers, uterine corpus endometrial carcinoma (UCEC) contained the largest number of lactylation-altering SNVs (20,428), followed by skin cutaneous melanoma (SKCM; 15,360) and colon adenocarcinoma (COAD; 7,291). The proportion of lactylation-altering SNVs varied across cancer types, ranging from 7.77% in pheochromocytoma and paraganglioma (PCPG) to 15.71% in SKCM. BLCA, READ, and CESC also showed relatively high proportions of predicted lactylation-altering SNVs (15.33%, 14.11%, and 14.02%, respectively). (Supplementary Figure 5A) Classification by positional relationship and predicted direction indicated that direct effects predominated over proximal effects. Direct lactylation gains constituted the largest category, followed by direct lactylation losses, whereas proximal gain and loss events were less frequent. Overall, predicted lactylation-gain events outnumbered lactylation-loss events, suggesting that cancer-associated missense SNVs more commonly confer a gain of predicted lactylation potential than a loss.(Supplementary Figure 5B) Within the 15 disease categories displayed for pathogenic variants, 211 of 2,923 pathogenic variants (7.22%) were predicted to alter lysine lactylation. Hereditary cancer-predisposing syndrome contained the largest absolute number of lactylation-altering pathogenic variants (34 of 555, 6.13%), followed by cardiovascular phenotype (19 of 313, 6.07%) and inborn genetic diseases (18 of 345, 5.22%). The highest proportions were observed for autosomal recessive limb-girdle muscular dystrophy type 2A (10 of 80, 12.50%) and PTEN hamartoma tumor syndrome (12 of 107, 11.21%).(Supplementary Figure 5C) These findings indicate that a subset of disease-associated pathogenic variants may contribute to disease mechanisms through predicted perturbation of lysine lactylation.

Furthermore, lactylated sequences exhibited distinct local amino acid preferences and sequence motifs, as described above (Figure 2J), suggesting that sequence context contains informative determinants associated with lactylation. We therefore reasoned that AttentionKla could capture relationships between local sequence patterns and lactylation propensity. Although the model assigns binary labels using a probability threshold of 0.5, its continuous prediction scores provide a relative measure of sequence-associated lactylation propensity rather than merely a binary classification outcome.

To investigate the sequence positions emphasized by the model, we extracted the attention matrices from the final Transformer layer and calculated the attention weights from the [CLS] token to each amino acid position by averaging across all attention heads. The resulting position-specific attention weights were subsequently aggregated across the selected sequences and fold models to generate an integrated mean attention profile. This attention-based positional salience profile represents the relative emphasis assigned by AttentionKla to each position during prediction and highlights candidate flanking positions that may contribute to the sequence context associated with lactylation (Figure 5B).

Consequently, mutating amino acids at these key positions—guided by in silico changes in model prediction values—could allow for the targeted modulation of lysine lactylation levels by leveraging the latent information learned by the deep learning model. Specifically, we selected SUMO2-K11la and ALDOA-K147la for experimental validation using site-specific lactylation antibodies^41^. The reported writers of these sites, AARS1 and p300, respectively, represent mechanistically distinct classes of lactyltransferases. We performed in silico mutagenesis of flanking residues at both sites and selected four variants predicted to have little effect, four predicted to increase lactylation, and four predicted to decrease it. In cells, mutation of the selected flanking residues produced concordant changes in SUMO2-K11la and ALDOA-K147la signals (Figure 5C-F): 6 of 8 variants predicted to increase lactylation increased the corresponding site-specific signal, whereas 4 of 8 variants predicted to decrease it reduced the signal. This directional asymmetry may reflect biological buffering and assay limitations: endogenous lactylation and delactylation activities can preserve residual site occupancy, while antibody-based measurements may have unequal dynamic ranges for gains and losses. When analyzed separately, model-predicted changes were positively correlated with experimentally measured changes in site-specific lactylation for SUMO2-K11la (Pearson r = 0.531, P = 0.0617, n = 13) and ALDOA-K147la (Pearson r = 0.646, P = 0.0172, n = 13). The pooled analysis across both proteins was also positive (Pearson r = 0.570, P = 0.0024, n = 26; Figure 5G). Notably, endogenous protein Kla signals could not be completely excluded during densitometric analysis, which may have upwardly biased the overall gray-scale values and reduced the quantitative precision of the estimated lactylation changes. This background contribution may therefore have introduced noise into the correlation analyses.

Although mutating flanking sequences may introduce new confounding variables, and the tool’s current success rate and generalizability require further investigation, this precise modulation of lactylation sites—achieved by extracting attention mechanisms from deep learning methods—provides a novel tool for investigating lactylation mechanisms and accurately controlling targeted modification levels.

## Discussion

By collecting lysine lactylation proteomics data, we constructed CLEAR-Lactyl, the largest human lactylation dataset to date, with stringent homology removal and false-negative control. Based on this dataset, we trained the AttentionKla model by fine-tuning ESM-2 with LoRA, achieving performance superior to existing tools. Notably, CLEAR-Lactyl includes 12.8% N-/C-terminally truncated sequences from all collected positive sites, which were padded with the pre-trained model’s <PAD> token, and negative samples containing <PAD> were supplemented accordingly. This design reduces concerns about information loss in atypical sequences; in performance benchmarking, the model achieved extremely high prediction accuracy on a pure truncated-sequence (only-<PAD>) testing set, with no sacrifice in performance on conventional sequences. This captures important functional signals and directly expands the practical utility of the model. Moreover, the unprecedentedly large scale of the lactylation site dataset lowers the false-negative rate during negative sample selection and demonstrates the potential of scaling laws. Nevertheless, constrained by the burden of data collection, we were unable to perform analysis starting from raw mass spectrometry data; consequently, the dataset is subject to the varying filtering criteria of the original publications, which may introduce certain biases. Despite this, AttentionKla still shows performance advantages on external testing sets spanning different disease types and even different species. Beyond powering AttentionKla, CLEAR-Lactyl represents a valuable resource for the broader research community, with the potential to support diverse investigations. Although such applications are beyond the scope of this study, CLEAR-Lactyl, AttentionKla, and proteome-wide predictions of lactylation propensity across human lysine sites, together with in silico mutagenesis outputs, are publicly available to support further investigation.

AttentionKla adopts ESM2-650M with LoRA, striking a balance between training cost and performance. ESM2, pre-trained on massive unlabeled sequences, has implicitly learned the “evolutionary grammar” of proteins as well as local physicochemical properties (e.g., amino acid charge and hydrophobicity). For lactylation—a “long-tail” modification with relatively limited positive data and species-specific variation—traditional CNN/RNN architectures are prone to overfitting, whereas based on pre-trained model, AttentionKla demonstrates powerful feature extraction and cross-species generalization^42^. Using AttentionKla, we generated a landscape of human lysine lactylation, in which 41% of residues emerged as potential sites, thereby defining the theoretical regulatory space of lactylation. Notably, the proportion of these potential sites that have been experimentally identified remains low, which may reflect the fact that many sites are modified only in specific spatiotemporal contexts or upon stress. Meanwhile, although the proportion of predicted potential lactylation sites is similar to that reported for other lysine PTMs, the number of high-confidence (FPR<5%) potential sites is markedly less than other reported lysine PTMs, possibly reflecting the true abundance of lactylation.

Analysis of the CLEAR-Lactyl dataset revealed an enrichment of lysine residues in the vicinity of positively labeled lactylation sites. On this basis, we hypothesized that lysine-rich local sequence contexts may promote lactylation by influencing substrate recognition, local electrostatic interactions, or conformational compatibility with lactyltransferases. One possible mechanism is that positively charged flanking residues interact favorably with acidic regions within the substrate-binding interface of a writer enzyme; however, this possibility remains to be established through structural and biochemical studies. More broadly, this hypothesis is consistent with the principle that the local sequence context contributes to substrate selection in several other post-translational modifications, including acetylation and ubiquitination.

This sequence-context preference further suggested that residues surrounding the central lysine may influence its lactylation propensity. Attention-based positional profiling, followed by in silico saturation mutagenesis, was therefore used to prioritize candidate flanking residues for experimental evaluation. Mutational analyses of SUMO2 and ALDOA provided proof-of-concept evidence that altering selected neighboring residues can modulate lactylation at the corresponding central sites. Together, these results indicate that AttentionKla captures interpretable sequence associations rather than functioning solely as a black-box classifier. Importantly, the attention weights should be interpreted as identifying sequence positions emphasized by the model and associated with its predictions, rather than as direct evidence that these residues causally determine lactylation.

Although this strategy successfully guided the modulation of two experimentally examined lactylation sites under distinct writer-enzyme contexts, its overall success rate and generalizability remain to be determined through broader validation across proteins, sites, cell types, and enzymatic contexts. The potential effects of the introduced mutations on protein structure, stability, localization, function, and other post-translational modifications at or near the target site must also be carefully considered. At present, this framework is therefore best regarded as a candidate-prioritization strategy that can reduce the experimental search space by directing mutagenesis toward a limited set of model-prioritized residues. More generally, it expands the investigation of lactylation regulation beyond the modified lysine itself to the contribution of its surrounding sequence context and provides a potential route for the targeted modulation of site-specific lactylation.

One limitation of this study arises from the nature of the training data. We used mass spectrometry data for model training, which reflects steady-state cumulative abundance and therefore cannot distinguish whether a modification arises from the activity of a synthetase versus a de-modification enzyme, nor can it capture the influence of dynamic factors such as intracellular lactate concentration or the spatiotemporal expression patterns of enzymes. Future work should incorporate time-series data and possibly other dynamic variables for training high-precision predictive models. Furthermore, although ESM2, the foundation of AttentionKla, has some implicit capacity for predicting spatial structure, a pure sequence-based model still struggles to directly capture steric hindrance and the effects of distal amino acid folding in three-dimensional space. Some lysines may be inaccessible to lactylation because distal residues shape folded conformations that bury or sterically occlude the lysine side chain. Integrating structural information from tools such as AlphaFold for multimodal representation is expected to improve prediction accuracy and spatial context awareness.^43^However, how to derive robust structural features and integrate them efficiently into model training without introducing excessive computational burden or information leakage remains an open challenge.

Despite these limitations, by constructing CLEAR-lactyl dataset, characterizing lactylation features, and training the AttentionKla model validated across diverse settings, we have achieved high-accuracy lactylation prediction and developed a strategy for modulating lysine lactylation levels through mutation of surrounding residues. Although lactylation levels can be controlled by K-to-R mutation or the PylRS approach, K-to-R substitution renders the target lysine non-lactylatable, whereas PylRS-based approaches can impose a defined modification state; both may therefore obscure the physiological consequences of dynamic and partial changes in lactylation occupancy. AttentionKla provides a selective flanking-residue mutagenesis approach, filling the gap in dynamically altering the modification level of a lactylated lysine. By extracting the model’s attention to local residues to guide site-directed mutagenesis, AttentionKla has transitioned from a “passive predictor” to an “active sequence designer”. By revealing the ability of flanking residues to drive lactylation, AttentionKla provides a framework that could advance our understanding of the relationship between lysine lactylation and human disease.

## Methods

### Dataset Construction

From publicly available human lactylation mass spectrometry (MS) data, we collected human lactylation sites identified by publicly available lactyl-proteomics studies. Only sites reported by the original studies at a false discovery rate (FDR) of <1% were retained. No restrictions were placed on sample origin to maximize site coverage. Centered on each identified lactylated lysine (K) residue, the sequence was extended by 25 amino acids towards both the N- and C-termini to extract a 51-aa sequence fragment. If a site was located near the protein terminus, resulting in fewer than 25 flanking amino acids, <PAD> tokens were used for padding. The CD-HIT program was employed for de-duplication and homology reduction of all sequence fragments, using a 90% sequence identity threshold to eliminate highly homologous relationships and retain representative sequences within each cluster. Following this process, the obtained site collection constituted the positive site standard set, comprising 16,604 lactylation sites.

For all protein involved in this positive set, we extracted the lysine residues that had not been identified as lactylated by any collected public data. For each such lysine, a 51-aa fragment (25 aa on each side) was similarly extracted to form the raw negative candidate pool. Applying the same criteria used for the positive samples, the raw negative candidate pool was subjected to de-duplication and homology reduction using CD-HIT with a 90% threshold to obtain a purified negative candidate pool. This step removed sequence redundancy and highly homologous sequences within the pool, while further reducing the probability of including undiscovered but true lactylation sites into the negative pool. From the purified negative candidate pool, an exact equal number of site sequences as the positive set (i.e., 16,604 sites) was randomly sampled to form the temporary negative set. The positive and negative sites meeting these criteria were merged to construct the CLEAR-Lactyl dataset, totaling 33,208 site entries.

### Motif Identification

To identify enriched sequence patterns flanking the lactylation sites, we performed a window-aligned recursive motif growth analysis between the positive sequences and the potential negative sequence sets. Local windows of 21 aa in length, anchored at the central site, were extracted. Only positive sequences without <PAD> tokens were retained for motif discovery, while the negative sequences were utilized for background amino acid frequency estimation. Background frequencies were calculated in a position-specific manner, where the occurrence frequency of each amino acid at every position within the negative sequence set served as the null distribution reference for subsequent enrichment tests.

Motif discovery was accomplished using a recursive greedy growth strategy. Each search iteration began with an initial pattern where the central lysine was fixed, and all other positions were designated as wildcards. Subsequently, the enrichment level was evaluated site-by-site and amino-acid-by-amino-acid within the current positive subset. Using the background frequency at the corresponding position in the negative set as the expected value, a one-sided binomial test was employed to screen for significantly overrepresented residues. At each step, the site-amino acid combination with the lowest *P*-value below a pre-set threshold was added to the current pattern, and the positive sequence subset was updated to include only sequences satisfying the new constraint. The current motif growth terminated when no additional sites met the significance threshold or when the number of remaining supporting sequences fell below a minimum threshold. The derived motif was recorded as a position-specific pattern, and the matched sequences were removed from the positive sequence set before continuing to the next search iteration until the minimum supporting sequence limit or the pre-set maximum number of motifs was reached. This method can progressively extract multiple relatively independent motifs.

### Structural interaction prediction and active-site pocket definition for lactyltransferases

Protein sequences of human AARS1, AARS2, EP300, GCN5, HBO1, TIP60, and KAT8 were retrieved from the UniProt database (https://www.uniprot.org/). These lactyltransferases, together with five virtual lactylation substrate sequences composed of the most frequent amino acid at each position within the 51-amino-acid window of positive sequences and five composed of the least frequent amino acid, were subjected to structural interaction prediction using AlphaFold3 (https://alphafoldserver.com/). A comprehensive analysis of residue-specific interactions between the proteins was performed, and all structural figures were generated using the open-source PyMOL molecular graphics system (version 3.2.0a, Schrödinger, LLC). For definition of the active-site pocket, the following residues were used: AARS1, 46, 77, 216, 239, 241; AARS2, 110, 128, 210, 240–242, 265, 269; EP300, 1398–1400, 1410, 1411, 1457, 1462, 1466, 1467; GCN5, 575, 579–581, 586–592, 617; HBO1, 475–477, 483–488, 512, 521, 508; TIP60, 370–372, 377–383, 407, 416, 403; and KAT8, 317, 319, 325, 326, 327, 329, 330, 354, 363, 408, 432, 350. Active-site pocket binding was defined as the presence of ≥3 residue-level contacts with interatomic distances <3 Å between the virtual lactylation substrate sequence and the corresponding enzyme pocket during non-polar interaction analysis in PyMOL; the shortest distance was recorded as the representative value for that enzyme– substrate pair.

### Enrichment Analysis

Functional enrichment analysis was performed on the Clear-Lactyl protein set using the Enrichr implementation in GSEApy. Gene symbols were converted to uppercase and queried against the GO Biological Process 2021, GO Cellular Component 2021, GO Molecular Function 2021, and KEGG 2021 Human gene set libraries, with Homo sapiens specified as the reference organism. Enriched terms were identified using the Enrichr statistical framework, and significance was assessed based on the adjusted P value, with terms passing adjusted P < 0.05 considered significant.

### Model Training

Training, validation, and testing data contained positive and negative samples. Each sample was a peptide sequence, with the lactylation site positioned at the center and flanking sequences of equal length (25 aa) on both sides. The positive samples were defined as sites with MS/MS evidence in CLEAR-Lactyl. For negative samples, we first retrieved all proteins from the CLEAR-Lactyl positive samples and comprehensively collected lysine sites on these protein sequences that lacked MS/MS validation to form the Potential Negative dataset. Subsequently, random sampling was employed to extract an equal number of negative sites from this dataset to match the positive sites, thereby preventing imbalanced training.

All peptide sequences were randomly split into three distinct sets: 80% for training, 10% for validation during training (representing the 9-fold cross-validation split), and 10% for internal independent testing (internal testing set). To further validate the generalization capability of AttentionKla, we constructed a new external independent testing set comprising a total of 4,426 site entries. This set included 2,213 de-duplicated human positive sites acquired from an independent study, and an equal number of negative sites sampled from the Potential Negative pool in the same manner. These testing sets, untouched during training and validation, were used to evaluate the model’s final performance.

In addition, we constructed a separate imbalanced external test set to assess model performance under a shifted class distribution. This set contained 1,391 positive sites with no overlap with the training data. For each protein contributing a positive site, the remaining lysine residues were considered candidate negatives; any sites overlapping with the training data were excluded before negative sampling. Negative sites were sampled to obtain a positive-to-negative ratio of 1:3. A matched balanced subset was generated by retaining all positive sites and randomly sampling one third of the negative sites.

Furthermore, to verify AttentionKla’s performance on data from other species, we selected lactylation MS data from four species with varying evolutionary distances from humans: *Mouse*, *Toxoplasma*, *Candida*, and *Salmonella*. Positive and negative samples were generated using the identical methodology (to enable simultaneous comparison between AttentionKla and the other three baseline models, only sequences without <PAD> were selected). This ultimately formed pan-species independent testing sets for *Mouse* (3,007), *Toxoplasma* (1,803), *Candida* (17,583), and *Salmonella* (438).

### Optuna Hyperparameter Optimization

To determine the optimal model configuration, we utilized the Optuna framework to perform Bayesian optimization on key hyperparameters for LoRA fine-tuning, executing a total of 50 search trials. The search space included learning rate (0.0009–0.0012, log scale), LoRA rank *r* (8 or 16), LoRA scaling factor α (16 or 32, constrained by α ≥ *r*), and training epochs (10–14). Other hyperparameters were kept fixed, including LoRA dropout at 0.3, target modules assigned to the query, key, and value projection layers of the attention mechanism, a training batch size of 64, and a cosine learning rate scheduler. Each candidate hyperparameter combination was evaluated via 9-fold stratified cross-validation to ensure consistent positive-to-negative sample ratios across all folds. For each fold, the model was initialized from the ESM2 pre-trained weights, and an early stopping strategy based on the validation set AUROC (with a patience of 2 epochs) was applied. The objective function for each trial was defined as the arithmetic mean of the AUROCs across the 9 validation folds. The hyperparameter combination yielding the highest mean AUROC was selected as the formal training configuration for AttentionKla.

### Benchmarking of Lactylation Predictions

To evaluate the performance of AttentionKla for Lactylation site prediction, we compared AttentionKla with three published tools, including AutoKla, DeepKla and AutoKla. For all public tools, we used the pretrained models made available by the original developers in the comparison.Among them, only AutoKla has the ability to handle sequence involving <PAD>. Therefore, the performance of Autokla was evaluated using the independent testing data (one outside and one inside) involving <PAD>, while DeepKla and PbertKla was evaluated using the independent testing data without <PAD> and compared with AttentionKla-nopad.

### Proteome-wide predictions for human

The FASTA file containing human protein sequences was downloaded from UniPort (accessed on 27 March 2026) and used as input to our deep learning models for lactylation site prediction. A probability score cutoff of 0.5 was applied to identify positive Lactylation predictions. Among these positive predictions, we further identified those with a probability score exceeding the threshold corresponding to a 5% false positive rate (FPR), as determined from the prediction results on the testing data.

### Identifying Variant-Induced Lactylation Changes

Pathogenic variants were obtained from ClinVar (https://www.ncbi.nlm.nih.gov/clinvar/, downloaded 21 May 2026). The Ensembl Variant Effect Predictor (v102.0, https://useast.ensembl.org/info/docs/tools/vep/index.html) was utilized for variant annotation. When variants could be mapped to multiple transcripts of the same gene, only the variant mapped to the canonical protein was retained, to eliminate redundancy. Somatic mutations were downloaded from the TCGA data portals (https://portal.gdc.cancer.gov/, accessed 17 May 2026). The Ensembl Variant Effect Predictor (VEP, release 113) was applied to annotate the functional consequence of all somatic mutations.. Only missense variants were used for downstream analysis. For each variant, 15-base peptides flanking the variant site were extracted from both reference and variant proteins as the input of Attention-Kla. A delta score cutoff of 0.5 was used to determine PTM-altering variants.

### Identifying Variant-Induced Lactylation Changes

Pathogenic variants were retrieved from ClinVar (downloaded May 21, 2026) and functionally annotated using the Ensembl Variant Effect Predictor (VEP, v102.0). When a variant was assigned to multiple transcripts of the same gene, only the consequence annotated on the Ensembl canonical transcript was retained to avoid redundant representation. Somatic single-nucleotide variants (SNVs) were obtained from the Genomic Data Commons (GDC) Data Portal for The Cancer Genome Atlas (TCGA) cohorts (accessed May 17, 2026) and annotated using VEP release 113. Only missense SNVs with an unambiguous protein-level consequence were retained for downstream analysis.

For each missense SNV, reference and variant protein sequences were generated. Candidate lysine lactylation sites located within seven amino-acid residues upstream or downstream of the mutated residue were identified. For each candidate lysine, sequence windows centered on the lysine residue and conforming to the input requirements of Attention-Kla were extracted from both the reference and variant proteins and used to predict lactylation probabilities. For direct effects in which a lysine residue was created or abolished by the variant, the lactylation probability of the sequence lacking the lysine residue was assigned a value of zero. The effect of each variant was quantified as ΔP=Pmu-Pwt, where Pmu and Pwt denote the predicted lactylation probabilities of the variant and reference sequences, respectively. Variants with ΔP ≥0.5 ΔP ≥0.5 were classified as lactylation-altering variants. Positive and negative Δ P values were interpreted as predicted lactylation gain and loss, respectively. Effects were defined as direct when the variant occurred at the candidate lysine site and proximal when the candidate lysine was located 1–7 residues from the variant position.

### Attention Weight Extraction and Positional Importance Analysis

To explore the biological basis of the model’s decisions, we performed attention analysis on the 200 positive sequences with the highest predicted probabilities in the testing set. For each sequence, it was fed into the 9 AttentionKla fold models independently. The attention matrices from the final Transformer layer were extracted, and the attention scores of the [CLS] token toward each sequence position were obtained by averaging across all attention heads. By accumulating and averaging the attention scores across all sequences and all fold models, we derived the integrated mean attention score for each position. This served as a quantitative indicator of the relative importance assigned to each position by the model, thereby illustrating the attention features of AttentionKla toward lactylation sequences.

### *In Silico* Positive Saturation Mutagenesis by AttentionKla

To evaluate the impact of the sequence context surrounding the target lactylation site on its lactylation propensity, we performed an *in silico* saturation mutagenesis analysis on the local sequence window centered around the target site. Specifically, for the 20 adjacent residues at relative positions -10 to -1 and +1 to +10 flanking the central site, we systematically introduced 20 standard natural amino acid substitutions, thereby constructing a single-point saturation mutant library. The central site itself was excluded from substitution to specifically evaluate the regulatory role of the neighboring sequence context. Notably, for input sequences containing <PAD>, we continued to treat <PAD> as a weighted token, ensuring the model’s capacity to comprehend sequence edge features was properly utilized. All candidate mutant sequences were input into AttentionKla for prediction. Using the predicted value of the wild-type sequence as a reference, we calculated the relative impact of each single-point substitution on the central site’s lactylation propensity. Ultimately, we ranked all substitutions based on this relative effect, defining the mutations with the strongest positive impact as positive saturation mutagenesis hits, and those with the strongest negative impact as negative regulatory candidates.

### Cell lines

Human cell lines of HEK-293T were obtained from the Chinese Academy of Sciences Cell Bank and authenticated via Short Tandem Repeat (STR) analysis. The cells were incubated in a humidified environment at 37 °C with 5% CO2 in DMEM high glucose medium (KeyGEN BioTECH, Nanjing, China), supplemented with 10% fetal bovine serum (EveryGreen, TIANHANG Biotechnology Co., Ltd, Huzhou, China), 100 μ/mL penicillin, and 0.1 mg/mL streptomycin (Beyotime, Shanghai, China).

To monitor potential Mycoplasma contamination, cells were routinely screened every two months through PCR detection. For this purpose, antibiotic-free culture supernatant was harvested following 7 days of cell growth. DNA was then isolated and cleaned up using silica-gel columns (TIANGEN, China). PCR amplification was carried out with hot-start Taq DNA polymerase following manufacturer’s instructions. The amplified products were electrophoresed on 1.3% agarose-TAE gels stained with ethidium bromide and analysed under UV light. A distinct band ranging from 515 to 525 bp would indicate Mycoplasma contamination. Only cell cultures confirmed to be Mycoplasma-free were utilized for further experimental procedures.

### Transient Transfection of Overexpression Plasmids

P53, SUMO2, ALDOA and their mutant sequences were codon-optimized and synthesized by GENEWIZ (Suzhou, China), and then cloned into integration-ready pcDNA3.1(+)-HA vectors. Plasmids were transfected into the HEK-293T cell line using the PEI 40000 reagent (YEASEN, #40816ES). The culture medium was replaced 8 hours post-transfection, and cells were harvested 72 hours post-transfection, followed by subjection to immunoblotting or immunoprecipitation.

### Immunoblot

Cells were lysed in RIPA buffer (MeilunBio, Dalian, China) containing a protease inhibitor and two phosphatase inhibitor cocktails (TargetMol), and maintained on ice for 10 min. After sonication, protein concentrations were determined using a BCA assay kit (YEASEN). Proteins were separated by SDS-PAGE gel (ShareBio, Shanghai, China) and then transferred to PVDF membranes (Millipore, Burlington, USA). Following a one-hour blocking step with 5% non-fat milk at room temperature, membranes were incubated with primary antibodies overnight at 4 °C and then exposed to HRP-conjugated secondary antibodies (1:2500, Beyotime) for 1 h at room temperature. Protein bands were detected using Enhanced Chemiluminescent Reagent (New Cell & Molecular Biotech, Suzhou, China). Details of the primary antibodies were provided as follows.

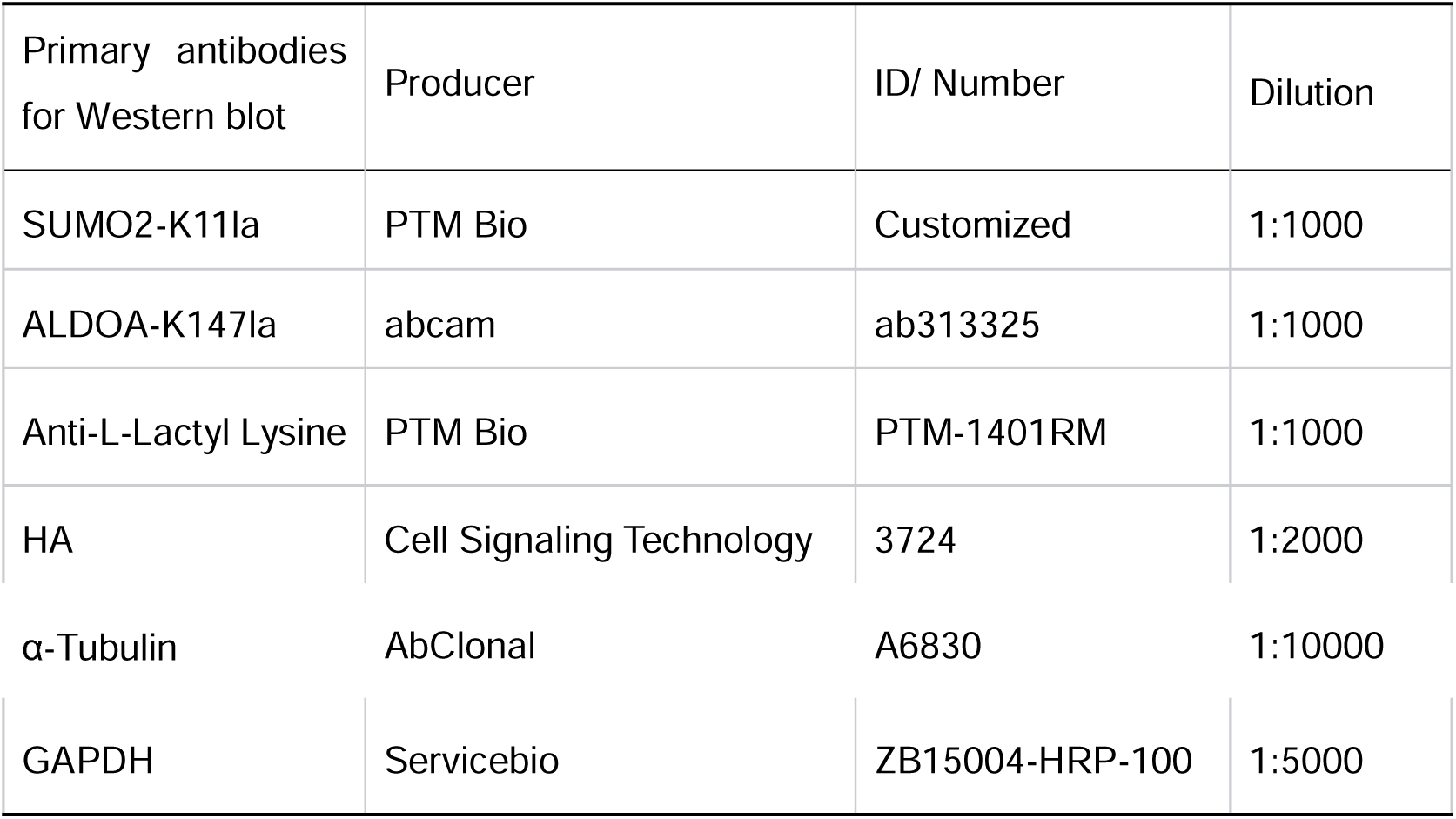

### Lactylation Modification Detection

Protein A/G magnetic beads (Thermo Fisher Scientific) were incubated with the indicated antibodies overnight at 4 °C and subsequently crosslinked with DSS. The beads were washed with NP-40 Lysis Buffer three times. Cell lysates were incubated with the crosslinked beads for a further overnight at 4 °C. Immunoprecipitated complexes were collected, washed, and then subjected to proteomics profiling or boiled with loading buffer in a dry bath heater for western blot detection. To investigate the lactylation level of transfected mutant p53, HA-p53 was immunoprecipitated utilizing an anti-HA antibody (1:50, #3724, Cell Signaling Technology, Danvers, USA) and then subjected to immunoblotting.

### Training Environment

Training was conducted on a server equipped with a single NVIDIA RTX 4090 24GB GPU, an Intel Xeon Gold 6430 CPU, and 120 GB of system memory, utilizing NVMe SSDs for storage. The GPU driver version was 560.35.03. The experimental environment was based on PyTorch 2.5.1, Python 3.12, CUDA 12.4, and cuDNN 9.1.0, with all code executed under the Ubuntu 22.04.5 LTS operating system.

## Supplementary information

Supplement Table: Source Data Supplement Figure: Supplement Figure1-4

## Code availability

The AttentionKla online prediction website is available at https://zhengdali-attentionkla.hf.space/. The model weights and associated code for AttentionKla are open-sourced at https://huggingface.co/ZhengdaLi/AttentionKla and https://github.com/lee190632/AttentionKla. The CLEAR-Lactyl and human whole-proteome lysine prediction results generated in this study are provided in the Supplementary Tables accompanying this article.

## Author contributions

Zhengda Li and Youcheng Huang performed dataset curation, model development, and formal analysis. Zhengda Li drafted the manuscript. Zhengda Li, Youcheng Huang, and Guangyao Shan performed the wet-lab experiments and experimental validation. Di Zuo, Dejun Zeng, Yuqiang Du, Junhe Zhang, and Xiliang Wang contributed to the bioinformatic analyses. Liang Chen, Hong Fan, and Guangyu Yao supervised the study and revised the manuscript. All authors reviewed and approved the final manuscript.

---

**Figure S1.**
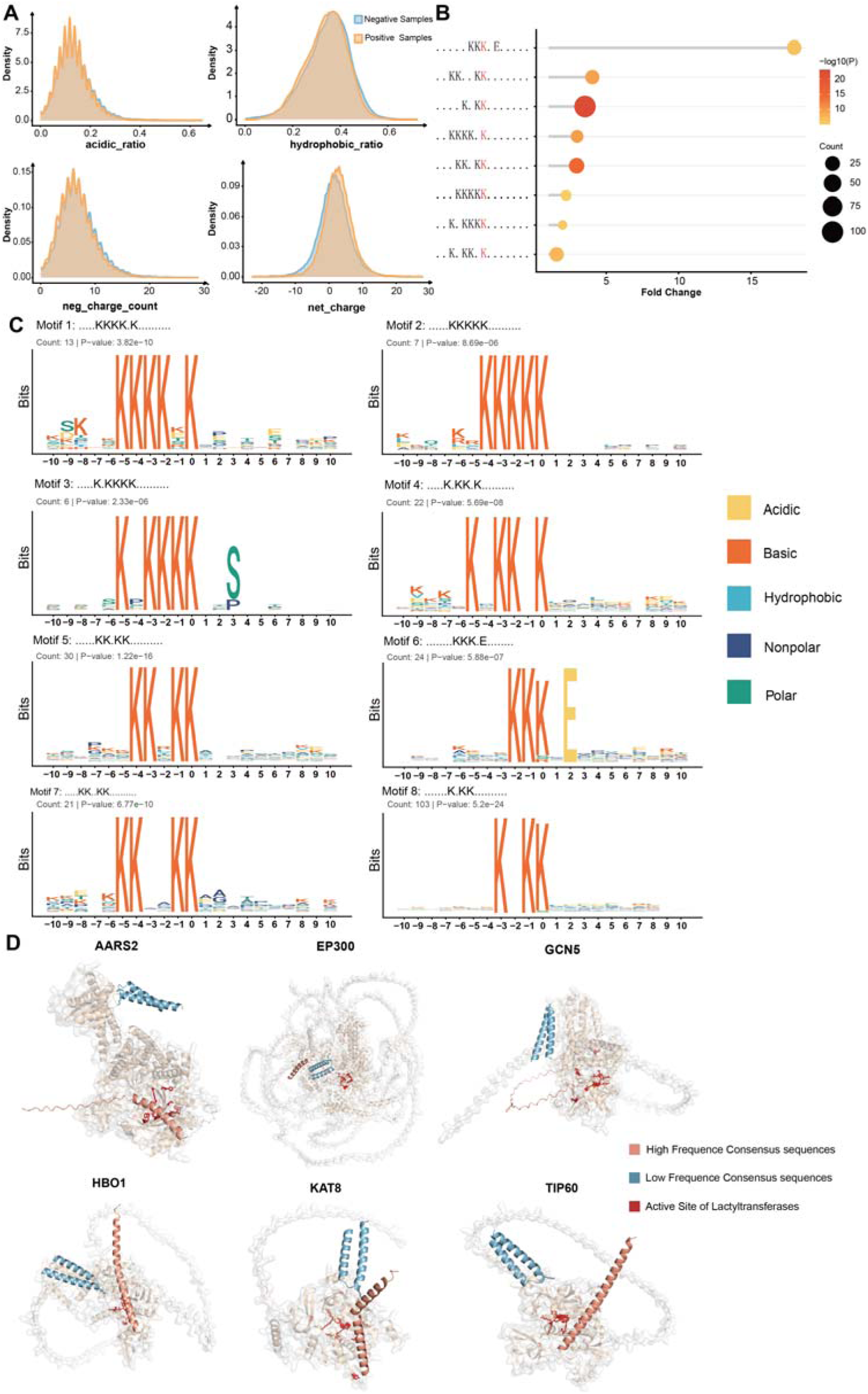
Supplementary analysis of sequence physicochemical properties and AlphaFold3 structural predictions. **(A)** Comparison of Relative Solvent Accessibility (RSA) scores between positive and negative lactylation sequences. **(B-C)** Motif discovery results for sequences extending 10 amino acids upstream and downstream of the positive lactylation sites, identifying key lysine-enriched motifs. **(D)** Extended AlphaFold3 prediction results assessing the interactions between consensus sequences and various lactylation enzymes, including both acetyltransferases and alanyl-tRNA synthetases.

**Figure S2.**
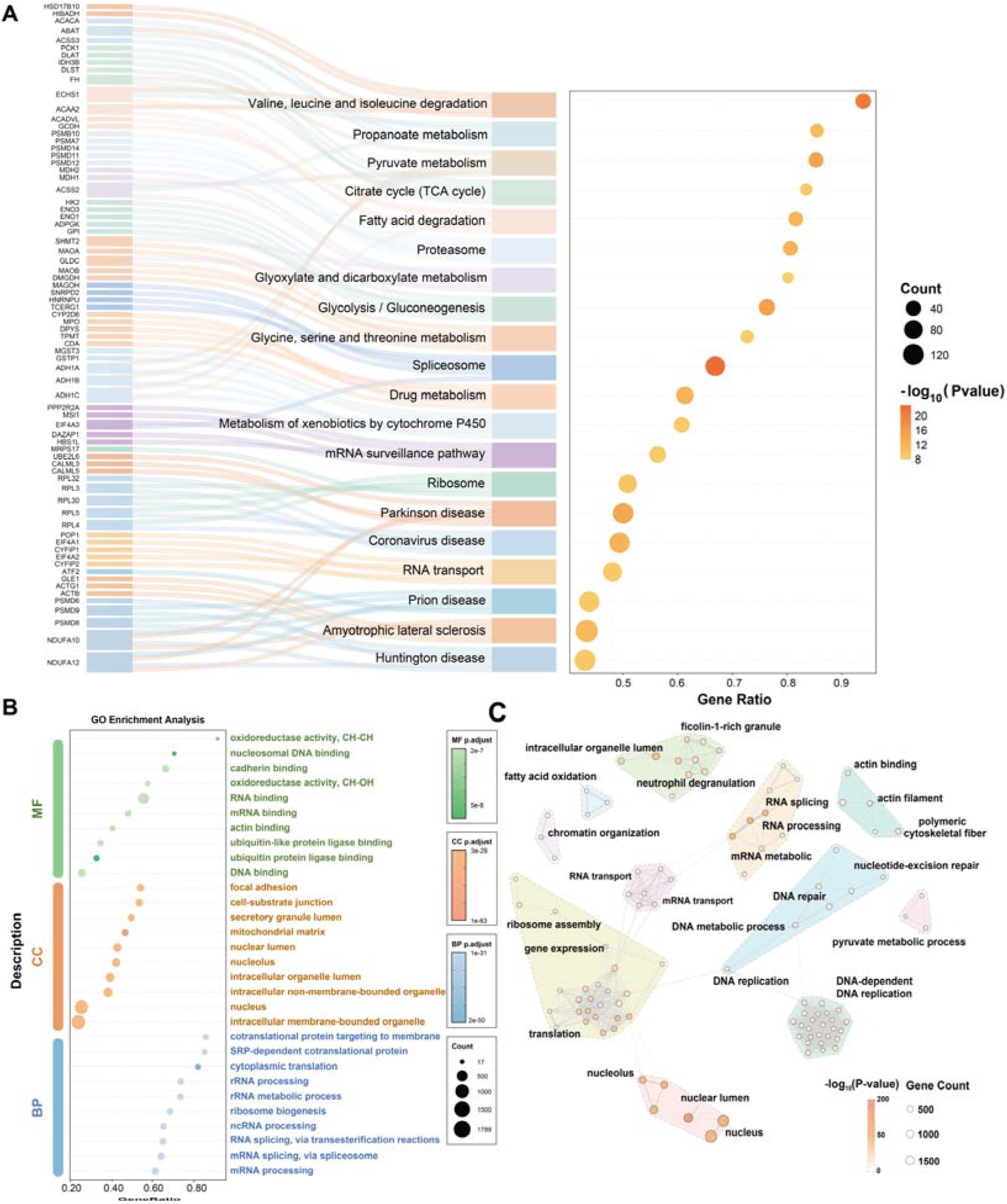
Functional landscape and enrichment network analysis of the human lactylome. **(A)** KEGG pathway enrichment and Sankey network analysis of the 4,934 lactylated proteins from the CLEAR-Lactyl dataset, highlighting the enrichment in classic metabolic pathways, core cellular complexes (spliceosome, ribosome, proteasome), and neurodegenerative diseases. **(B)** Gene Ontology (GO) enrichment analysis illustrating the top significantly enriched terms across Molecular Function (MF), Cellular Component (CC), and Biological Process (BP). **(C)** Functional enrichment network displaying highly interactive modules regulated by lactylation, including hubs for RNA splicing/processing, ribosome assembly/translation, DNA replication/repair, and fatty acid oxidation.

**Figure S3.**
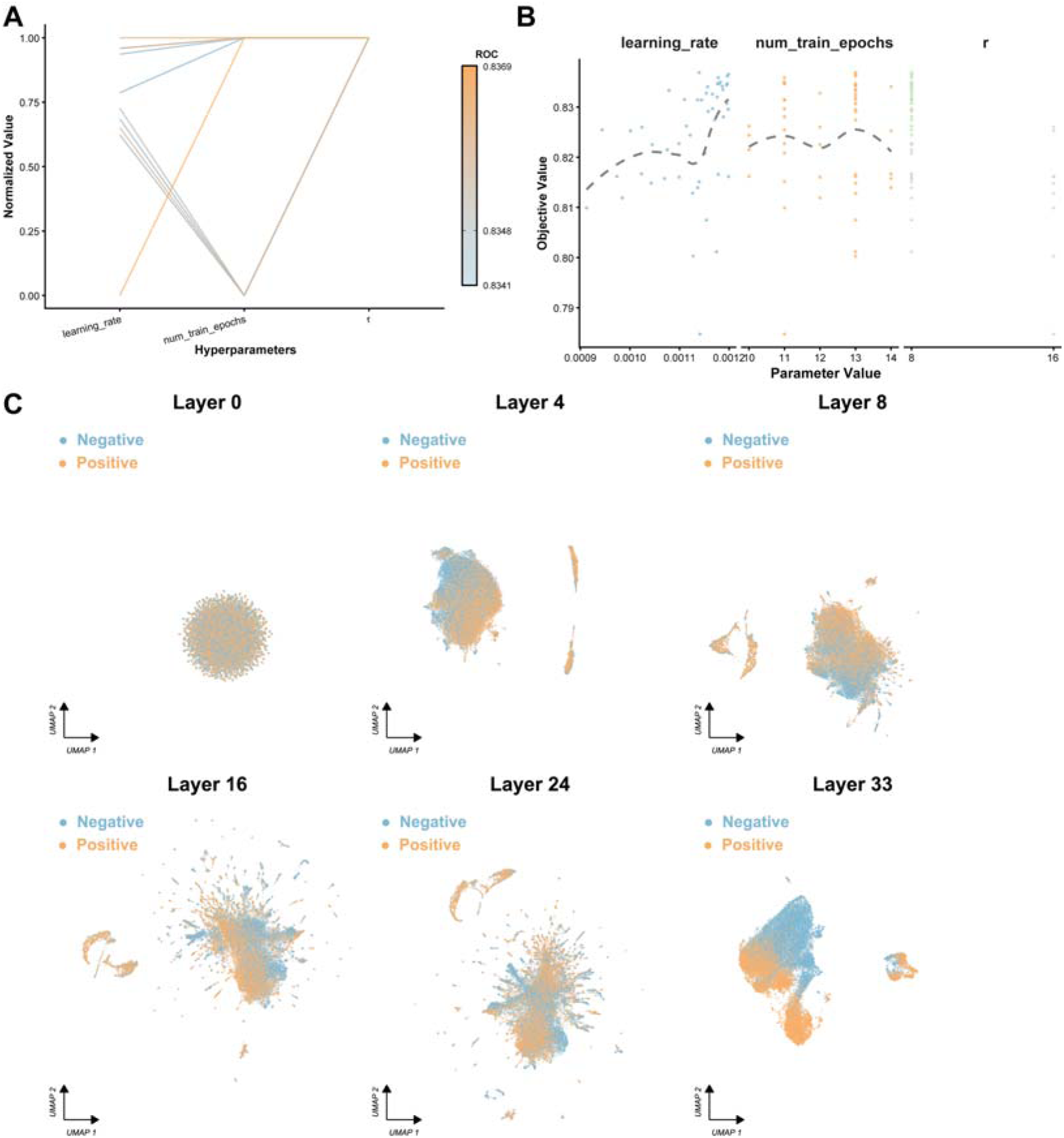
Hyperparameter optimization and feature representation of AttentionKla. **(A-B)** Optuna-based hyperparameter search landscapes for fine-tuning the ESM2-650M model, evaluating combinations of learning rate, LoRA rank, and alpha over 50 trials. **(C)** UMAP (Uniform Manifold Approximation and Projection) visualization of sequence features extracted from different layers of AttentionKla, illustrating the progressive separation of positive and negative lactylation sites throughout the network depth.

**Supplementary Figure S4.**
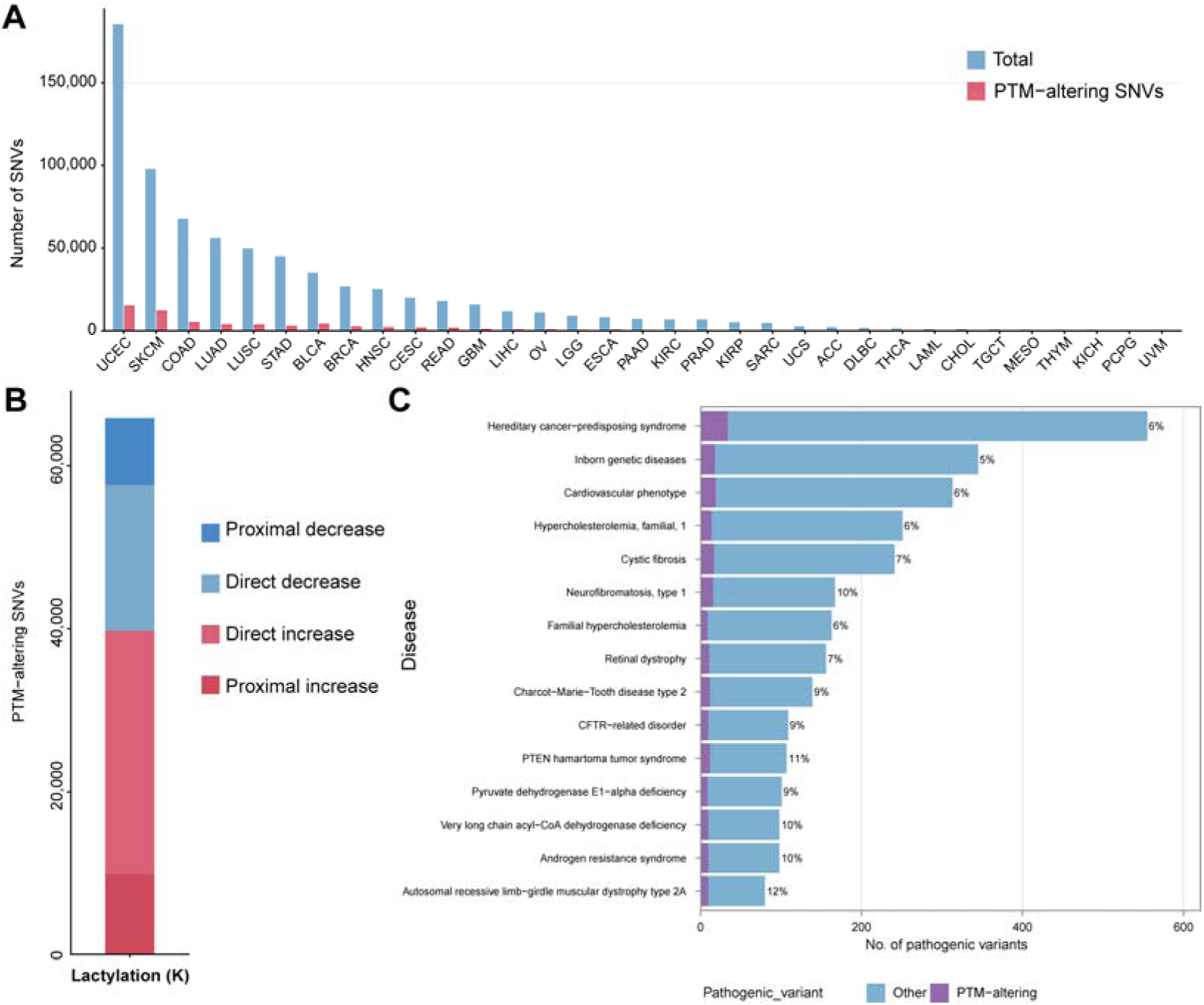
Ablation studies and baseline re-training results evaluated on the internal testing set. **(A)** Performance comparison of different fine-tuning strategies evaluated on the internal testing set. **(B)** Impact of foundation model parameter scales on AttentionKla performance on the internal set. **(C)** Performance evaluation on the imbalanced and balanced testing set.**(D-F)** Extended evaluation metrics for the AutoKla, DeepKla, and PBertKla models evaluated on the internal testing set after being re-trained with the CLEAR-Lactyl dataset. **(G-I)** Extended performance comparisons of AttentionKla and AttentionKla-nopad across specific internal sub-testing datasets: **(G)** mixed sets, **(H)** only-no <PAD>, and **(I)** only-<PAD>.

**Figure S5.**
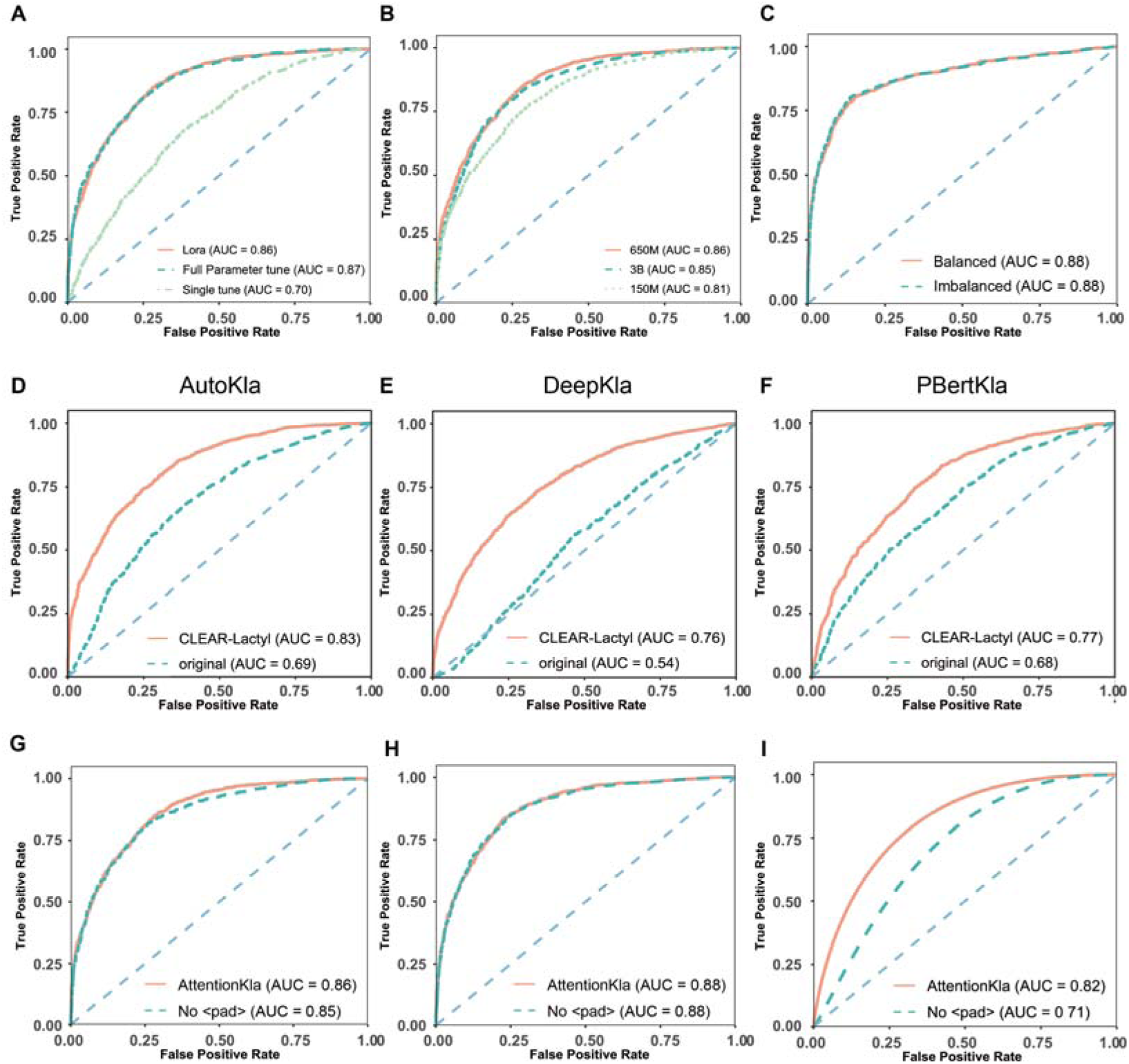
Pan-cancer and disease-associated landscape of predicted variant-induced lactylation alterations. **(A)** Numbers of total somatic missense SNVs and predicted lactylation-altering SNVs across 33 TCGA cancer types. Cancer types are ordered by total SNV number. **(B)** Distribution of predicted lactylation-altering SNVs according to positional relationship and effect direction. Direct effects denote variants occurring at the candidate lysine site, whereas proximal effects denote variants affecting lysine sites located within 1–7 amino-acid residues of the variant. Positive and negative changes in predicted lactylation probability are denoted as increase and decrease, respectively. **(C)** Distribution of ClinVar pathogenic variants across the 15 displayed disease categories. Stacked bars show predicted lactylation-altering variants and other pathogenic variants; percentages indicate the proportion of lactylation-altering variants within each disease category. A variant was classified as lactylation-altering when the absolute difference between the predicted lactylation probabilities of the variant and reference sequences was at least 0.5.

